# A novel integrated extraction protocol for multi-omic studies in heavily degraded samples

**DOI:** 10.1101/2023.12.15.571815

**Authors:** Byron Boggi, Jack Sharpen, George Taylor, Konstantina Drosou

## Abstract

The combination of multi-omic techniques, e.g. genomics, transcriptomics, proteomics, metabolomics and epigenomics has revolutionised studies in medical research. These are employed to support biomarker discovery, better understand molecular pathways and identify novel drug targets. Despite concerted efforts in integrating omic datasets, there is an absence for the integration of all four biomolecules in a single extraction protocol. Here, we demonstrate for the first time a novel, minimally destructive integrated protocol for the simultaneous extraction of artificially degraded DNA, proteins, lipids and metabolites from pig brain samples. We used an MTBE-based approach to separate lipids and metabolites, followed by subsequent isolation of DNA and proteins. We have validated this protocol against standalone extraction protocols and show comparable or higher yield of all four biomolecules. This integrated protocol is key towards facilitating preservation of irreplaceable samples while promoting downstream analyses and successful data integration by removing bias from univariate dataset noise and varied distribution characteristics.

---

Over the last decade, multi-omic techniques have been in the forefront of biological sciences. The analytical power integrating multiple datasets has facilitated major advances in medicine, microbiology, modern human disease studies^1,2^ ecology and agriculture^3^. The purpose of multi-omic studies is to provide a deeper understanding of the interplay between multiple layers of biological regulation and in particular to support biomarker discovery, predict novel drug targets, increase the diagnostic power for health and improve disease prognosis^4^. Despite the potential of machine learning technologies, multi-omic data integration faces many challenges such as data normalisation and compatibility, clustering, functional characterisation and visualisation which confound their analyses^5,6^. An added level of complexity is that multi-omics datasets are derived from blending data from individual workflows, different processing platforms, and intra-sample biomolecule preservation.

In heavily degraded remains, i.e. ancient, forensic, archival or even clinical Formalin-Fixed-Paraffin-Embedded (FFPE) samples, the situation is even more challenging as sample heterogeneity, limited DNA quantity and fragmentation, degradation and contamination, post-mortem damage and chemical modifications can all pose challenges and further bias in downstream analyses which complicates data integration^7,8^. For these reasons, multi-omic analyses and data integration in the ancient DNA (aDNA) field have been massively overlooked and to this day most research focusses either on genomics, or proteomics, and most often individually, rather than through a combined approach. A major drawback is that archaeological samples are rare and irreplaceable so there is a need to increase the amount of data that can be derived from a single biopsy. Integration on the workflow level, i.e., the extraction of two or more biomolecules from a single sample/biopsy, has however been limited, and the few examples where it has been attempted in the literature have concentrated largely on the combination of DNA and protein for example from dental remains^9^ and dental calculus in particular^10^. Furthermore, in the former case, the efficacy of the combined protocol was not compared to similar, individual protocols, making it difficult to assess the effectiveness of the method. The latter study reported a nearly 50% decrease in endogenous DNA recovered when compared to a DNA only protocol likely due to the method requiring long incubation times (72h) with no nuclease protection. Previous studies have shown that an optimised silica based spin column can effectively co-extract DNA, RNA and protein^11^ and commercial kits have been available for a number of years (AllPrep DNA/RNA/Protein Mini Kit - Qiagen). Such kits could likely be optimised for aDNA and peptide recovery in a similar fashion to other proprietary chemistry^12^. One of the main practical challenges to co-extract aDNA and peptides is the long incubation time during extraction and fragile nature of aDNA in particular, whereby inhibition of nucleases is essential to preserve as much genomic content as possible. As with most DNA extraction protocols, proteinases are the gold standard. However, such treatments are not amenable to proteomic workflows for already degraded peptides in bottom-up workflows. Hence, inactivation of nucleases could be achieved with a strong denaturant, such as methanol, where loss of structure will effectively inhibit nuclease activity. Furthermore, such denaturants will lead to the precipitation of genomic and proteomic contents of the extracted sample and partially purify the sample and facilitate the recovery of polar and non-polar small molecules. As such, solvent mixtures (chlorofolm:methanol)^13,14^, have been adopted as the gold standard for lipids and metabolites for over 70 years and are widely used for the co-extraction of protein^15^.

Another popular biphasic organic extraction protocol for lipids and metabolites, adopts Methyl-tert-butyl-ether (MTBE) as a safer alternative to chloroform and allows faster and cleaner recovery of a broad range of lipid classes^16^. As with chloroform:methanol this protocol has been shown to effectively co-extract proteomic content and provide a comprehensive view of cell signalling mechanisms^17^.

However, a protocol that includes DNA, lipids, proteins, and metabolites does not currently exist in published literature for ancient, forensic or clinical samples. Here we demonstrate a novel, fully integrated extraction protocol implemented on artificially degraded porcine brains that is suitable for degraded samples. We designed our protocol with three considerations:

First, we consider that biomolecular analyses of archaeological remains involve destructive sampling and require a significant amount of initial template for separate extraction methods. In addition, such analyses are often complemented by radiocarbon dating and stable isotope analysis for which additional sample material is required. Ancient soft tissues, such as hair, skin, muscle and brain, have rarely been considered for biomolecular and biomedical research, because oftentimes, these are part of an intact, mummified individual, and these types of remains are subject to different sampling considerations than skeletal remains. Therefore, by combining the extraction of ancient DNA, proteins, and lipids from a single biopsy we significantly reduce the amount of template material, thus preserving a finite and irreplaceable resource, and facilitate greater depth of analysis that would otherwise be prohibited due to the requirement for destructive sampling.

Second, of all the soft tissues preserved in the archaeological record, brain is one of the most overlooked tissues as its recovery was previously considered to be rare phenomenon. This tendency to refer to it as rare, has caused a paucity into ancient brain tissue analyses, but also studies in brain preservation, and degradation mechanisms^18^. However, to date there are over 200 independent published reports about preserved brain specimens from a wide range of depositional environments and around a thousand more reports about brain specimens dating back to the 17^th^ century in which the brain appears in various states of preservation^19–24^. Moreover, forensic studies in brain tissue have found that DNA is relatively resistant to putrefaction and in contrast to bone, the evidence suggests a certain degree of protection against DNA degradation^25,26^. Such studies have been supported by Serrulla^26^ et. al. who recovered 45 well preserved brains from a Spanish civil war mass grave. In this case the wet conditions at the level of the burials have led to poor skeletal preservation but exceptional soft tissue preservation including brains^27^. Despite brain tissues having been recovered from both forensic and archaeological environments, biomolecular research from brain tissue is extremely limited with only a handful of examples^28,29^. Therefore, the necessity of having an optimised protocol in place to allow the extraction of biomolecules from soft tissues that are resistant to degradation is highlighted.

Third, in aDNA studies, mineralised matrices such as bones and teeth are preferred, largely because early studies showed that in comparison to skeletal remains, DNA derived from soft tissues are significantly more fragmented, the tissues had a low endogenous DNA content and they only survive under certain climatic conditions in the archaeological record^30^. As a result, most methods upstream of high throughput DNA sequencing (HTS) have been optimised for mineralised samples. However, soft tissues can be more informative than skeletal tissues, especially in palaeopathological studies as they carry different molecular information^31^. Brain samples can facilitate omics analyses with unprecedented resolution as neurodegenerative and psychiatric disorders are affected by a multitude of factors such as pathogen-driven selection, genetics and environmental factors the signatures of which can be detected with omics analyses, specifically through lipidomics and proteomics. In addition, brain samples can play a key role in elucidating evolutionary patterns as brain has been shaped over the course of time^32^. Other soft tissues such as muscle and skin can offer a more direct representation of an individual’s metabolic state including disease-related markers or evidence of exposure to toxins. Therefore, developing an extraction protocol that is optimised for soft tissues and facilitates multi-omics analyses is of paramount importance.

Therefore, we have devised an integrated two-step protocol of extracting lipids, metabolites, proteins and DNA. We combined a well-established solvent-based approach that facilitates co-extraction of lipids and metabolites through phase separation by centrifugation. This results into two layers which contain the non-polar lipids (top phase) and polar metabolites (lower phase), leaving denatured protein and DNA pelleted at the bottom of the tube. The second step involves the resuspension of DNA and protein, separating the DNA from the protein-tissue pellet with a final precipitation step, leaving the DNA soluble in the supernatant. The subsequent extracts are then ready to be prepared for downstream data acquisition.

## Results

### Artificial desiccation results

Four pig brain biopsies (SS1, SS2, SS3, SS4) from the cerebral cortex were obtained and each was split into two hemispheres. These were desiccated at different time-points and divided into four samples. A total amount of 2mg of cerebral cortex was collected from the frontal lobe to facilitate extraction using a) standalone (SA) protocols of DNA^12^, protein, lipid and metabolite^14^ extractions and b) our integrated (INTG) extraction protocol.

Morphological changes to the brain samples that were typical of artificial desiccation experiments were observed over the course of four timepoints. These included shrinkage of the cerebral cortex and decrease in weight due to loss of water content. On a molecular level we found that there is approximately 15% post-mortem modifications (PTMs) in proteins, of which 50% suggest deamidation. Upon further inspection, we found deamidation in proteins not associated with these modifications *in-vivo*, such as Syntaxin binding proteins and tubulin, whilst also identifying lysine acetylation at expected residues on Serine and arginine rich splicing factor 3 within both protocols. This demonstrates that our artificial desiccation protocol successfully mimics spontaneous mummification. Despite the limitations in obtaining quantitative estimation of DNA degradation using Bayesian computation we employed a qualitative approach to assess DNA sequence characteristics indicative of degradation. Through visual inspection, we observed signs of fragmentation, low base quality scores and a decrease in mean sequence length, all of which indicate damaged sequences suggestive of degradation.

### DNA results

DNA recovery showed the largest deviation when compared to the SA protocol from equivalent masses of brain tissue. All four samples showed a significant increase in yield which is evident from the amount of raw and mapped reads per chromosome (**Table 1**) as well as the number of single nucleotide polymorphisms (SNPs) recovered (**Table 2**).The sequencing results for both the INTG and SA protocols can be found in Table 1. In line with increased yields of DNA, the number of raw reads following sequencing was significantly higher from the integrated protocol compared to that of the dedicated DNA extractions where we have an approximate average of 27M reads compared to 16M reads respectively. However, the increased number of reads is to be expected given the higher DNA concentration recovered, but it should be noted that identical elution volumes were used to resuspend the DNA post extraction. Furthermore, for DNA samples processed by the dedicated protocol, on average >60% of reads aligned to the *Sus Scrofa* genome were removed during the duplicate removal stage of data pre-processing. This is indicative of a less-than-optimal sample preparation where the PCR is dominated by a low field of correctly formatted amplifiable library molecules.

**Table 1.** Mapping Statistics per library and mapped reads per chromosome. Each sample yielded between 7.1 and 52.7 million reads, with an average of 60% of the reads uniquely mapping to the Sus Scrofa genome. The samples consistently exhibited a substantial amount of endogenous content, ranging from 5.3% to 61.2%. The average fragment lengths varied between 55 and 69bp.

|  | SS1<br>SA | SS1<br>INTG | SS2<br>SA | SS2<br>INTG | SS3<br>SA | SS3<br>INTG | SS4<br>SA | SS4<br>INTG |
| --- | --- | --- | --- | --- | --- | --- | --- | --- |
| Raw reads | 7,003,531 | 36,467,461 | 21,619,544 | 13,386,446 | 21,761,393 | 40,849,208 | 15,540,794 | 52,678,198 |
| Mapped reads | 4,496,535 | 22,075,298 | 13,667,758 | 9,399,508 | 10,680,890 | 26,018,098 | 10,290,120 | 38,969,751 |
| Unique reads | **814,656** | 16,941,890 | **7,956,412** | **7,326,233** | 4,045,118 | 7,702,800 | **3,278,082** | 32,096,857 |
| Redundancy | 81.87% | 23.25% | 41.78% | 22.05% | 62.12% | 70.39% | 68.13% | 17.63% |
| μ read length | 60bp | 56bp | 55bp | 68bp% | 68bp | 62bp | 69bp |  |
| End. DNA | 11.6% | 46.5% | 36.8% | 54.7% | 18.6% | 18.8% | 21.0% | 60.9% |
| Chr1 | 85,904 | 133,158 | 803,696 | 752,368 | 427,869 | 808,989 | 345,566 | 3,233,544 |
| Chr2 | 49,126 | 1,662,529 | 501,769 | 471,091 | 245,168 | 479,456 | 207,966 | 2,155,310 |
| Chr3 | 46,128 | 1,088,957 | 483,881 | 447,903 | 229,480 | 451,855 | 196,438 | 2,073,002 |
| Chr4 | 43,105 | 1,059,710 | 414,411 | 384,060 | 212,955 | 408,760 | 175,313 | 1,703,578 |
| Chr5 | 34,394 | 879,573 | 346,039 | 322,773 | 171,922 | 331,634 | 145,233 | 1,466,299 |
| Chr6 | 59,542 | 745,787 | 627,085 | 573,367 | 294,955 | 583,271 | 253,859 | 2,713,065 |
| Chr7 | 40,976 | 1,390,611 | 411,008 | 372,069 | 203,856 | 392,386 | 169,593 | 1,686,410 |
| Chr8 | 43,164 | 878,092 | 399,120 | 378,586 | 216,363 | 406,545 | 173,414 | 1,604,841 |
| Chr9 | 45,355 | 833,367 | 429,374 | 397,536 | 221,642 | 423,235 | 180,345 | 1,737,267 |
| Chr10 | 23,742 | 899,091 | 235,310 | 213,352 | 119,332 | 225,803 | 97,399 | 949,637 |
| Chr11 | 25,114 | 503,701 | 238,235 | 227,187 | 124,018 | 236,179 | 101,802 | 977,115 |
| Chr12 | 23,523 | 509,816 | 261,811 | 240,418 | 112,750 | 229,708 | 102,836 | 1,197,572 |
| Chr13 | 64,133 | 608,040 | 600,899 | 559,136 | 320,681 | 603,339 | 258,840 | 2,380,203 |
| Chr14 | 47,232 | 1,232,094 | 475,958 | 431,745 | 235,702 | 457,671 | 196,264 | 1,963,887 |
| Chr15 | 44,087 | 1,017,433 | 408,414 | 382,245 | 218,323 | 413,062 | 176,667 | 1,643,580 |
| Chr16 | 25,458 | 845,872 | 254,141 | 229,567 | 125,801 | 237,558 | 103,267 | 963,285 |
| Chr17 | 22,050 | 494,678 | 230,570 | 208,237 | 108,455 | 212,637 | 92,550 | 968,453 |
| Chr18 | 19,182 | 506,110 | 191,335 | 176,804 | 95,352 | 184,820 | 78,909 | 801,665 |
| ChrX | 36,067 | 414,699 | 345,218 | 325,376 | 187,258 | 342,860 | 74,495 | 703,636 |
| ChrY | 268 | 724,315 | 2,323 | 2,455 | 1,196 | 2,532 | 10,224 | 89,596 |
| MT | 15,749 | 6,483 | 61,921 | 23,159 | 42,193 | 57,335 | 41,859 | 171,291 |

**Table 2.** Variant calling per sample, per chromosome. SA: Standalone Protocol, INTG: Integrated Protocol.

| Chromosome | RefSeq | SS1<br>SA | SS1<br>INTG | SS2<br>SA | SS2<br>INTG | SS3<br>SA | SS3<br>INTG | SS4<br>SA | SS4<br>INTG |
| --- | --- | --- | --- | --- | --- | --- | --- | --- | --- |
| Chr 1 | NC_010443.5 | 0 | 3,506 | 87 | 274 | 10 | 73 | 15 | 22,595 |
| Chr 2 | NC_010444.4 | 0 | 4,784 | 126 | 431 | 5 | 73 | 14 | 24,844 |
| Chr 3 | NC_010445.4 | 2 | 5,298 | 134 | 314 | 21 | 117 | 12 | 28,906 |
| Chr 4 | NC_010446.5 | 4 | 3,104 | 66 | 239 | 27 | 69 | 22 | 16,470 |
| Chr 5 | NC_010447.5 | 6 | 3,230 | 224 | 400 | 43 | 152 | 100 | 19,317 |
| Chr 6 | NC_010448.4 | 1 | 5,391 | 149 | 441 | 17 | 106 | 16 | 19,648 |
| Chr 7 | NC_010449.5 | 2 | 2,372 | 75 | 179 | 9 | 51 | 11 | 19,899 |
| Chr 8 | NC_010450.4 | 2 | 1,793 | 150 | 268 | 24 | 98 | 45 | 14,880 |
| Chr 9 | NC_010451.4 | 1 | 2,392 | 172 | 353 | 30 | 107 | 25 | 17,676 |
| Chr10 | NC_010452.4 | 4 | 2,060 | 118 | 203 | 21 | 142 | 23 | 12,010 |
| Chr 11 | NC_010453.5 | 0 | 1,638 | 27 | 125 | 2 | 50 | 0 | 10,906 |
| Chr 12 | NC_010454.4 | 0 | 4,977 | 112 | 360 | 18 | 60 | 11 | 23,295 |
| Chr 13 | NC_010455.5 | 1 | 1,879 | 93 | 171 | 12 | 54 | 18 | 13,213 |
| Chr 14 | NC_010456.5 | 0 | 2,915 | 77 | 247 | 6 | 46 | 8 | 22,075 |
| Chr 15 | NC_010457.5 | 8 | 2,292 | 100 | 209 | 17 | 55 | 20 | 12,205 |
| Chr16 | NC_010458.4 | 12 | 1,280 | 121 | 202 | 78 | 125 | 62 | 7,703 |
| Chr 17 | NC_010459.5 | 1 | 2,501 | 78 | 186 | 6 | 40 | 19 | 13,331 |
| Chr18 | NC_010460.4 | 0 | 1,564 | 37 | 117 | 1 | 29 | 3 | 9,915 |
| Chr X | NC_010461.5 | 3 | 814 | 29 | 75 | 0 | 18 | 0 | 1,440 |
| ChrY | NC_010462.3 | 0 | 30 | 2 | 3 | 0 | 1 | 509 | 2,009 |
| MtDNA | NC_000845.1 | 173 | 182 | 177 | 174 | 19 | 20 | 20 | 1,440 |

The most notable difference between the two protocols was observed during variant calling. The integrated protocol yielded a much higher number of SNPs across all chromosomes. Manual inspection of regions that were mutually covered by aligned sequencing reads exhibited a greater depth of coverage favouring the integrated protocol (**Table 2**). We further analysed samples SS2 (SA) / SS2 (INTG) which have largely comparable numbers of aligned reads (in total and per chromosome) and found a significantly higher number of SNPs obtained from the integrated protocol. We compared the read to SNP ratio of the unfiltered VCF files and the percentage of SNPs passing filtering (MQ<30, DP>5, QUAL30, QD<2.0, SQR>3.0, FS>60.0, MQRankSum<-12.5, ReadPosRankSum<-8.0) was much higher from the INTG protocol. For example, sample SS2 (SA) has a higher number of aligned reads overall and following read duplicate removes compared to its equivalent SS2 (INTG) (**Table 1**). Despite the higher number of aligned reads the number of SNPs passing quality filtering are more than double for the INTG protocol. Furthermore, the percentage of reads passing quality filtering was much higher (X% vs Y%) for the INTG approach. Due to the differences in the number of raw reads per library, the remaining samples could not be directly compared. However, we normalised the data and analysed SNP-to-aligned read ratio, again finding that it was significantly higher in favour of the integrated protocol. Our results suggest that, when considered holistically, DNA preparation using the integrated protocol was more successful by comparison to the standalone approach throughout all stages, from sample preparation to computational processing.

### Protein results

Following shotgun proteomics, we validated peptide sequence matches (PSMs) of raw files through MaxQuant^33^ search using MSStats^34^ for processing and summarisation, before comparing SA with INTG protocols for both normalised peak intensities of total proteins discovered and categorising their general functions. Both protocols show that most proteins are related to neuronal and metabolic functions, which is expected within brain samples. Furthermore, proteins and isoforms specific to the brain and nervous system were the dominant class in each protocol respectively, showing a high degree of uniformity. Whilst there were more PSMs in the SA protocol than the INTG post processing, we found that INTG SS2 had significantly less proteins recovered and at lower peak intensities than the other 3 INTG samples, indicating a loss or disruption of sample in the process. In contrast, a greater overall recovery of individual protein abundance in SS1-SS3 samples such as with the ubiquitous tubulin, spectrin, alpha (axon actin-ring cytoskeleton) and syntaxin binding protein 1 (neurotransmitter release in synaptic vesicles), further validates the comparability between protocols.

### Lipid and metabolite results

The initial lipid spectral identifications showed that in general, the integrated protocol recovered more peaks across both Mass Spectrometric Modes than Bligh and Dyer^14^, with a significant increase within Positive Mode. The only sample which had significantly less recovery when using the multi-omic extraction protocol was SS2 Negative Mode lipids which further supports a loss of sample during the process through processing error. When examining the lipid family composition the protocols are highly comparable and consistent with current brain lipid literature; the predominance of glycerophospholipids in both modes is expected due to being the main constituents of neuronal cell membranes^35^ whilst the significance abundance of sphingolipids are commonly found within the brain due to their roles in neurogenesis and synaptogenesis^36^. These general observations of tissue specificity and lipid recovery are similarly reflected when discriminating into lipid classes), where sphingomyelin and its biosynthetic ceramide precursors, both highly enriched in oligodendrocytes and myelin, consist of the majority positive mode identifications, whereas the membrane-based phosphatidylethanolamine (PE) and its derivatives dominate the negative mode, whilst also elucidating cardiolipin, a hallmark of inner mitochondrial membrane that’s predicted to be abundant within the brain^37^.

Statistical analysis of small molecules and metabolites recovered and identified showed no significant difference between both Integrated and traditional protocols used, with comparative MS peak identifications found. Compounds specific to a range of metabolic pathways were ubiquitously found in all samples, not limited to oxidative phosphorylation (e.g. maleic acid, citric acid, ADP), protein and lipid metabolism, as well as neurotransmitters such as glycine, glutamate and GABA derivates in γ-aminobutyrate. As such, these results are comparable and are broadly consistent with previous comparisons between Bligh and Dyer^14^ and MTBE-based protocols^16,17^

## Discussion

This proof-of-concept paper demonstrates for the first time the successful implementation of a novel multi-omic integrated protocol for the simultaneous extraction of DNA, proteins, metabolites and lipids from a single degraded sample. This approach increases the acquisition of omics data from a single sample whilst reducing time, associated costs, and more importantly sampling requirements (which is of paramount importance, when finite or even archival and clinical FFPE samples are processed).

Four porcine brain samples were artificially degraded for up to three months at 37°C and subsequently sub-samples were processed using well-established standalone protocols for the individual extraction of DNA^12^, proteins, lipids and metabolites^14^. Equivalent sub-samples of the same four samples were also processed using our integrated protocol. Comparisons between the integrated and standalone protocols demonstrate that our multi-omic protocol is either comparable or superior to the single-omic protocols. Both protocols generated a substantial number of raw reads, with the SA protocol producing between 6 and 21 million reads, while the INTG protocol produced between 13 and 52 million reads. Discrepancies in the raw data could be attributed to differences in DNA concentration, pooling and sequencing bias. However, when we normalised the results to account for biases, we observed a consistently better results in favour of the integrated protocol for all samples (except for SS1, which exhibited more than twice the proportion of mapped reads compared to raw reads (**Table 1**), but this did not translate into higher rates of SNP calling). Additionally, when mapping to the *Sus Scrofa genome*, the samples processed with the INTG protocol showed a significant percentage of high-quality SNP calls per sample compared to the dedicated protocol (**Table 1**), even when the number of aligned reads was lower than the equivalent SA sample (**Tables 1, 2**). Aligned reads per chromosome generally showed comparable or better results between the two protocols, except for a few instances where the SA protocol exhibited more Y chromosomal reads (**Table 1**). In addition, there was increased depth of coverage which is of great significance for applications in ancient and forensic fields, as it decreases the need for additional target enrichment. Similarly to the DNA results, protein recovery was also comparable or higher in all four samples apart from sample SS2 which is likely a processing error. Organic molecule, metabolite and lipid recovery was consistently comparable among all samples with the INTG protocol recovering more identifications in both positive and negative modes during lipidomic analysis.

So far, biomolecular analyses of degraded brain samples have been extremely rare, with only two successful examples documented in the literature^27,28^. following singe-omic approaches. Additionally, one study produced dubious results^29^ due to the lack of adherence to strict anti-contamination criteria. With rare exceptions, the standardised and comprehensive analysis of ancient, forensic, mummified and other heavily degraded brain tissues has been notably lacking on a large scale. This gap signifies a vast reservoir of unexploited tissue resources that could significantly contribute to the exploration of brain evolution. As a consequence, contemporary research on human brain evolution^38–41^ has predominantly relied on modern samples, primarily focusing on the genomic level. Computational approaches, cross-species comparative genomics as well as population genetic-level approaches have been instrumental in exploring the diversity of the human genome. Notably, these studies have highlighted the role of Transposable Elements (TE) as influential drivers of human brain evolution^32^.

However, pathogenesis encompasses a complex interplay of various regulatory elements such as proteins, metabolites, lipids, genes and epigenetic features such as DNA methylation and histone post-translational modifications (PTMs). Considering brain is abundantly found in mummified remains and is resistant to degradation due to its high lipid content, multi-omics studies including both ancient and modern data, are critical in providing insights into the intricate dynamics underlying brain evolution such as genetic, epigenetic and biochemical factors. Furthermore, it is possible to unravel the mechanisms behind well-documented brain preservation in the archaeological record.

By successfully characterising biomolecules from degraded brain samples we systematically address the limitations posed by degraded soft tissues, thereby challenging preconceived notions regarding the applicability of such samples in biomolecular/biomedical research, and re-assess their value within the context of brain evolution and pathogenesis. In particular, the protocol’s ability to reduce processing time, material requirements but also biases associated with sample preservation variations and are in turn responsible for deviation of results and lack of authentication and replication, enhances its practical utility. Processing a single sample under the same protocol and storage conditions, has the potential to help with noise reduction and artefacts for downstream bioinformatic applications by facilitating top-down approaches to simultaneous integration and dimensionality reduction of both NGS and MS datasets, which will further improve robustness for subsequent machine learning methods. This quality is especially valuable in studies involving finite or archival samples, where obtaining meaningful data can be challenging.

Additionally, we underscore the brain’s significance as a model organism and lay the groundwork for future investigations into various tissues. The initial focus on brain tissues serves as a strategic starting point acknowledging the inherent challenges associated with these samples. As the protocol proves its efficacy in the brain, the potential to other tissues becomes evident, promising a versatile tool for multi-omic analyses in diverse biological contexts.

## Methods

### Integrated (multi-omic) extraction

The protocol was validated using the same four artificially desiccated samples (SS1, SS2, SS3, SS4) against dedicated well-established and widely used extraction protocols for DNA, protein, lipids and metabolites. Subsequent extracts were processed for individual downstream workflows and data acquisition performed via high throughput sequencing (HTS) for DNA and liquid chromatography with tandem mass spectrometry (LC-MS/MS) for protein, lipids and metabolites. For the integrated (INTG) and standalone (SA) protocol for DNA, vortexing and ultrasonication were avoided to prevent fragmentation of the DNA. This is important in this study as we want to ensure this protocol is suitable for forensic and archaeological studies and case work.

### Step 1: Organic molecule separation

For our INTG protocol, we opted for a relatively large volume of extract as we did with the standalone protocol^14^. The initial ratio of methyl-tetr-butyl ether (MTBE) and methanol (MeOH) (10:3 *v/v*) based lysis remains true to the original description^16^. However, we included 0.01% (butylated hydroxytoluene) BHT in our MTBE:MeOH mixture, to minimise oxidation of lipids. Furthermore, we incorporated the use of 0.1% ammonium acetate (AA) to induce phase separation^17^ whilst aiding precipitation of insoluble material including DNA and protein. Lipids and metabolites were first extracted using 4ml MTBE:MeOH in a 10:3^17^ ratio with a 60 minutes incubation step at 4°C with gentle agitation in a shaking incubator. Phase separation was induced with 770μL 0.1% AA (10:3:2.5 (*v/v/v*)) to aid precipitation of insoluble material, including protein and DNA. The upper layer, containing the non-polar lipids, and the lower layer, containing the polar metabolites, were then decanted into fresh glass tubes for drying and resuspension. Extracts were then dried under a gentle flow of nitrogen at 40oC and re-suspended in either 200μL MTBE containing 0.01% BHT (lipids) or 200μL of 80% aqueous CAN with 0.1% FA (metabolites) and submitted to the Bio-MS core facility at the University of Manchester for data acquisition.

### Step 2: DNA and Protein separation

The resultant pellets from the original extraction tubes were left to air dry for 5 - 10 minutes. The pellets were then resuspended in 500μl of 5% sodium dodecyl sulfate (SDS) based lysis buffer (pH8) and left to incubate at room temperature for 30 minutes with gentle agitation. Following centrifugation at 1500xg for 10 minutes to pellet any insoluble material, the supernatant was transferred to 2ml DNA Lobind tubes (Eppendorf) and 250μL AA (7.5M) was added for a final concentration of 2.5M. Following the addition of AA, samples were left to incubate at room temperature for 15 minutes and the protein was pelleted by centrifugation at 16,000xg for 10 minutes. The resultant supernatant, containing DNA, was then suspended in five volumes of spiked Qiagen PB buffer^12^ and purified by MinElute (Qiagen) with the addition of reservoirs for larger sample volumes^12^ followed by two rounds of elution with 30 μl elution buffer (Qiagen EB). Extracts were stored at -20°C until library preparation and sequencing at the Genomics Facility of the University of Manchester for data acquisition.

The remaining protein pellet was resuspended in 500μl of a protein lysis buffer consisting of 5% SDS, 50mM Tetraethylammonium bromide (TEAB) and LC-MS grade H^2^O, (PH 7.5). All samples were reduced with dithiothreitol (DTT), at a final concentration of 5mM, for 10 minutes at 60°C and allowed to cool to room temperature. Alkylation was facilitated with iodoacetamide (IAM) at a final concentration of 15mM with a 30 minute incubation at room temperature in the dark followed by quenching with DTT. Samples were then centrifuged at 14,000xg for 10 minutes to pellet insoluble debris and the resultant clarified lysate was transferred to a fresh protein LoBind tube (Eppendorf).

Lysates were quantified on a Qubit™ 4 fluorimeter, with the Protein Broad Range assay kit (Invitrogen). Trypsin digestion was performed using the S-Trap column digestion protocol^42^ (ProtiFi). All subsequent binding, washing and elution steps are followed by centrifugation at 4000xg for 2 minutes. Samples were diluted to 1mg/ml and 50μl (containing 50μg) and was acidified with 5μl of 12% aqueous phosphoric acid. The total 55μl of acidified sample was loaded onto the columns with 330μl S-Trap binding buffer (90% MeOH, 100mM TEAB, PH 7.1). One wash with MTBE, followed by four washes with S-Trap binding buffer was performed. Lyophilised trypsin was dissolved in 50mM TEAB and loaded onto the columns, (final protease: protein ratio of 1:10) and incubated for 1 hr at 47°C. Three elution steps were performed with 65μl of 50mM TEAB, 65μl of 0.1% aqueous Formic Acid (FA) and 30μl of 30% aqueous acetonitrile (ACN) containing 0.1% FA. Samples were further desalted using Oligo™ R3 reversed-phase resin (Thermo Scientific) and eluted twice in 50μl of 30% aqueous ACN containing 0.1% FA. Samples were dried by SpeedVac and submitted to the Bio-MS core facility, The University of Manchester, Manchester, UK for data acquisition.

Standalone Protocols were conducted following previously established protocols^12,14^.

## Acknowledgments

The authors extend their appreciation to the Genomic Technologies Core Facility, the BioMS Facility and the Computational Shared Facility (CSF) at the University of Manchester. For advice and additional training we thank Professor Anna Nicolaou. We extend our special thanks to Emeritus Professor of Biomolecular Archaeology Terence A. Brown for his advice and support.

